# Aging Differentially Affects Axonal Autophagosome Formation and Maturation

**DOI:** 10.1101/2022.08.27.505553

**Authors:** Heather Tsong, Erika Holzbaur, Andrea KH Stavoe

## Abstract

Misregulation of neuronal autophagy has been implicated in age-related neurodegenerative diseases including Parkinson’s disease and Huntington’s disease. We compared autophagosome formation and maturation in primary murine neurons during development and through aging to elucidate how aging affects neuronal autophagy. We observed an age-related decrease in the rate of formation of LC3B-positive autophagosomes leading to a significant decrease in the density of autophagosomes along the axon. Next, we assessed the maturation of autophagic vesicles and identified a surprising increase in their maturation in neurons from aged mice. While we did not detect notable changes in endolysosomal content in the distal axon during aging, we found that autophagic vesicles were transported more efficiently in neurons from adult mice than in neurons from young mice. This efficient transport of autophagic vesicles in both the distal and proximal axon is maintained in neurons from aged mice and indicates that aging alone does not impair transport nor negatively impact the later stages of autophagy. However, the pronounced deficit in autophagosome biogenesis in aged neurons may contribute to a decreased capacity to clear aggregated proteins or dysfunctional organelles and thus contribute to age-related degeneration.

## INTRODUCTION

Macroautophagy (hereafter autophagy) is a degradative process that is integral to maintaining cellular homeostasis of eukaryotic cells. Autophagy is especially critical in neurons, as neurons are post-mitotic, metabolically very active, and highly polarized cells with functional compartments such as synapses localized far away from the buffering capabilities of the cell soma. Furthermore, adult neurogenesis occurs infrequently and individual neurons must be retained throughout an animal’s lifespan. The importance of autophagy in sustaining neuronal function is demonstrated by early-onset neurodegeneration in mice with neuron-specific knockout of critical autophagy genes^1–3^. Additionally, dysregulation of autophagy has been implicated in age-related neurodegenerative diseases including Alzheimer’s disease (AD), Parkinson’s disease (PD), Huntington’s disease (HD), and Amyotrophic Lateral Sclerosis (ALS)^4–6^.

Autophagy is a complex but evolutionarily-conserved pathway originally delineated in yeast and predominately studied in yeast and mammalian cell culture in the context of acute stressors such as starvation^7–12^. The pathway begins with autophagosome formation – the de novo creation of a double membrane structure engulfing, in the case of bulk macroautophagy, nonspecific cytoplasmic cargo and organelle fragments. Over 40 autophagy-related genes (“ATG”) have been identified in mammals, the majority of which coordinate autophagosome biogenesis. The Initiation complex, the Nucleation complex, the Elongation complex, the ATG2 complex, and other individual ATG proteins work in concert to build and extend the double membrane structure. During biogenesis, the outer leaflet of the double membrane becomes decorated with phosphoethanolamine (PE)-conjugated microtubule-associated light chain 3B (LC3B) and its orthologs, members of the ATG8 family. LC3B and other ATG8 proteins are the only known proteins that mark autophagosomes both during and after formation, as the rest of the autophagy machinery dissociates from the organelle prior to closure of the autophagosome membrane to generate the hallmark double membrane structure. Following closure, PE-conjugated LC3B remains in the luminal-facing leaflet, while the PE-LC3B present on the outer, cytosolic face is cleaved off the autophagosome^13^.

Once autophagosome biogenesis is complete, the autophagosome fuses with a late endosome or lysosome to generate an autolysosome. The internal pH of the autolysosome drops as lysosomal V-type ATPases acidify the compartment, leading to the activation of degradative enzymes that breakdown internalized cargos to facilitate recycling of their components^14^.

In neurons, the stages of autophagy are spatially and temporally regulated. Autophagosome biogenesis predominately occurs in the distal axon^15–19^. Once formed, autophagosomes are transported along microtubules. Initially, autophagosomes exhibit bidirectional movement in the distal axon, enabling initial fusion with distal late endosomes and lysosomes^15^. Subsequently, immature autophagosomes and maturing autolysosomes, jointly termed autophagic vesicles (AVs), undergo processive retrograde movement to the cell soma directed by cytoplasmic dynein. Cargo degradation initiates en route, while the final delivery of AVs to the soma may enhance the recycling of component macromolecules into biosynthetic pathways to generate new proteins and organelles^1,15,18,20–26^.

Many studies have linked defects in autophagy to age-related neurodegenerative diseases^4–6^. Here, we examined autophagosome biogenesis, transport and maturation in murine dorsal root ganglia (DRG) neurons during development and early aging using live imaging approaches that provide high temporal and spatial resolution. DRG neurons are a powerful model, as these cells are one of the few neuronal types that can be cultured from any age mouse, enabling us to interrogate the autophagy pathway in neurons across relevant timescales^27^. We used this model to interrogate the specific steps involved in autophagosome biogenesis, maturation, and trafficking to better understand which steps in the pathway are altered during aging. Together, our results indicate that while autophagosome biogenesis slows significantly, most of the subsequent steps in the pathway are not adversely affected by aging, thus defining a specific target for possible therapeutic intervention to enhance autophagy in aging neurons.

## RESULTS

### Autophagosome Biogenesis and Transport Flux Decrease with Age

Misregulation of autophagy has been implicated in neurodegeneration. Since age is the most common risk factor for neurodegenerative disease^28^, we asked how each of the steps of basal autophagy might be affected by aging in murine neurons. Using GFP-LC3B transgenic mice^29^, we assessed the rate of autophagosome biogenesis in primary DRG neurons from three different ages of mice: 1-month-old young mice, 3-month-old young adult mice, and 16-17-month-old aged mice. First, we examined the recruitment of autophagy components required for autophagosome biogenesis, focusing on markers for the initiation complex (mCherry-ATG13)^30–32^ and the nucleation complex (Halo-ATG14)^33–35^. We used live-cell imaging to observe the recruitment of each marker to developing phagophores in the distal axon (Figure 1A). We did not observe any significant changes with age in the appearance of either mCh-ATG13 (Figure 1B) or Halo-ATG14 (Figure 1C) puncta, demonstrating that these early stages of autophagosome biogenesis are not altered during aging in DRG neurons. Additionally, we did not observe any significant changes in the protein levels of ATG13 and ATG14 in immunoblot of whole brain lysates (Figure S1A-B). In striking contrast and consistent with our previous results^27^, we observed a significant deficit in the rates of autophagosome formation corresponding to a 41% drop between neurons from young and aged mice (Figure 1D), consistent with the dysregulation of autophagosome biogenesis at the axonal terminal of DRG neurons during aging.

**Figure 1.**
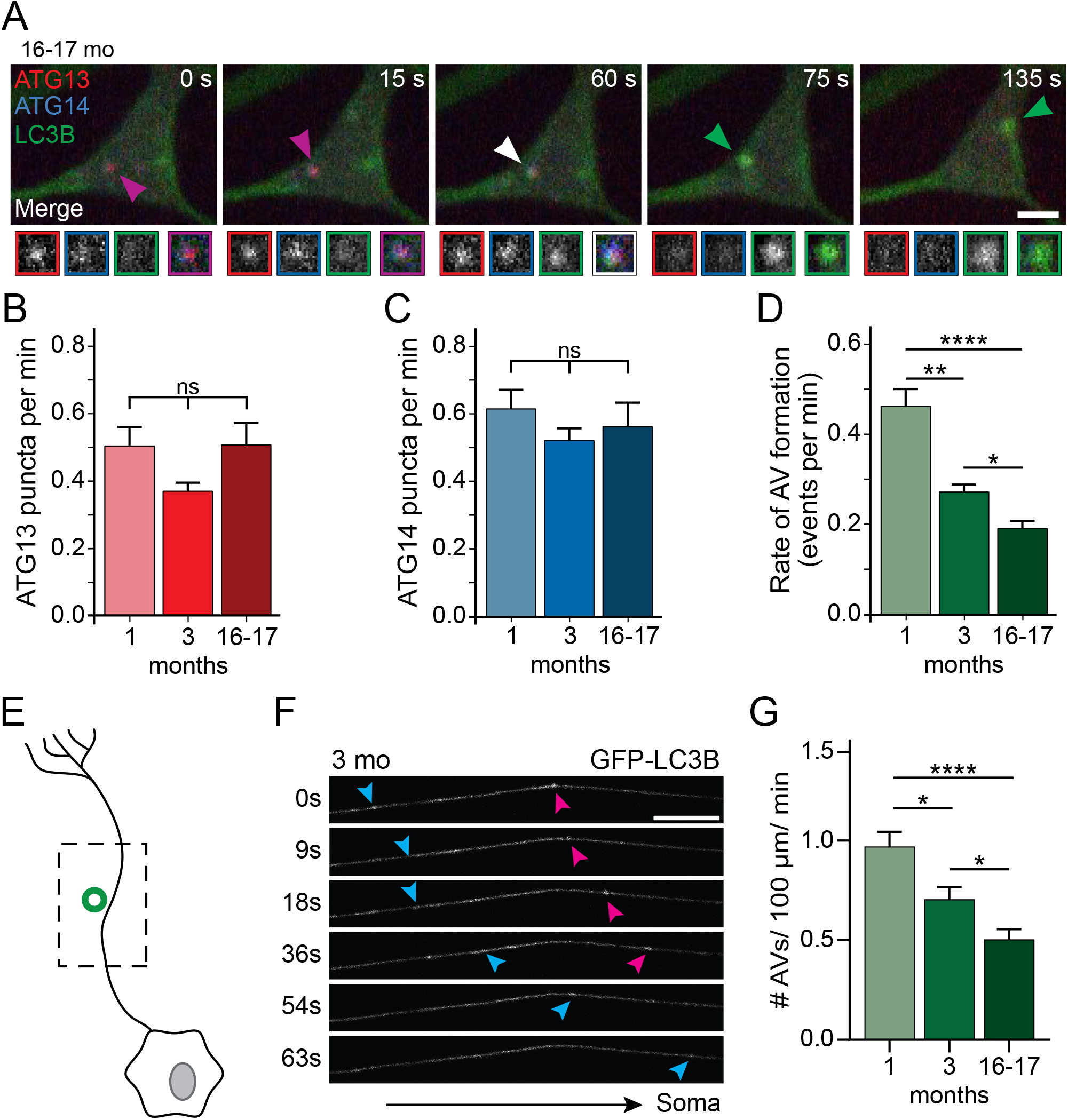
Distal autophagosome biogenesis and mid-axon autophagosome transport flux decrease with age. (A) Time series of merge micrographs of mCh-ATG13, Halo-ATG14, and GFP-LC3B from live cell imaging of the distal neurite of DRGs from aged mice depicting an autophagosome biogenesis event. Purple arrowheads denote colocalization of mCh-ATG13 and Halo-ATG14; white arrowhead denotes colocalization of mCh-ATG13, GFP-LC3B and Halo-ATG14; green arrowheads denote a GFP-LC3B-positive punctum from which mCh-ATG13 and Halo-ATG14 have dissociated. Magnified views of denoted puncta are shown below full micrograph; border color represents channel (left 3 grayscale boxes per timepoint) or colocalization state in merge (right box per timepoint). Retrograde is to the right. Scale bar, 2 μm. (B-C) Quantification of the rate of mCh-ATG13 (B) and Halo-ATG14 (C) puncta formation in live-cell imaging of DRG neurons from young, young adult, and aged mice (mean ± SEM; n≥ 18 neurons from three biological replicates). ns, not significant by Kruskal-Wallis test with Dunn’s multiple comparisons test. (D) Quantification of the rate of autophagic vesicle (AV) biogenesis (assayed by GFP-LC3B puncta formation per minute) in DRG neurons from young, young adult, and aged mice (mean ± SEM; n≥ 18 neurons from at least three biological replicates). ****p < 0.0001; **p < 0.005; *p < 0.05 by Kruskal-Wallis test with Dunn’s multiple comparisons test. (E) Cartoon of DRG neuron depicting time-lapse imaging of AVs in the mid-axon. (F) Representative time series of micrographs of GFP-LC3B from live-cell imaging in the mid-axon of DRGs from a young adult mouse. Colored arrowheads denote two individual GFP-positive puncta moving retrograde. Retrograde is to the right. Scale bar, 10 μm. (G) Quantification of the number of GFP-positive puncta detected per minute per 100 μm in the mid-axon of DRG neurons from young, young adult, and aged mice (mean ± SEM; n≥ 90 neurons from three biological replicates). *p < 0.05; ****p < 0.0001 by Kruskal-Wallis test with Dunn’s multiple comparisons test.

Once autophagosomes form in distal axon, they undergo retrograde trafficking toward the cell soma^15^. We asked if transport of autophagosomes in the axon changes with age. First, we assessed transport flux of autophagosomes in the mid-axon using live-cell imaging of the GFP-LC3B probe in primary DRG neurons (Figure 1E). We counted the number of GFP-LC3B puncta observable during a three-minute imaging window (Figure 1F) and normalized for the length of the axon segment within the micrograph frame. We detected a significant decrease in the transport flux of AVs with age corresponding to a 28% drop between neurons from young and young adult mice and 48% drop between neurons from young and aged mice (Figure 1G). Taken together, these data suggest that autophagosome biogenesis decreases in the distal axon with age leading to a significant decrease in the transport flux of autophagosomes in the mid-axon with age.

### Autophagosome maturation in the distal axon increases with age

The marked decrease in autophagosome biogenesis with age is one factor contributing to the age-related decrease in the transport flux of AVs in the mid axon. Other factors may also contribute, including alterations in the retrograde transport of AVs in the axon, driven by the molecular motors cytoplasmic dynein and kinesin along the microtubule cytoskeleton^15^. Fusion of autophagosomes with lysosomes and late endosomes, or the acidification of AVs following fusion may also be altered in aging.

The pH of autophagic vesicles directly affects the ability to detect GFP-LC3B-marked AVs. Since LC3B remains only inside the autophagosome after closure^13^, once the pH of the autolysosome falls below the pKa of GFP, the fluorophore is quenched. The pH-sensitive quenching of GFP-LC3B may have a significant impact on the interpretation of AV transport flux data in the mid axon. Thus, we subsequently monitored AV transport in the axon using a tandem mCherry-eGFP-LC3B probe^36^. Since the pKa of mCherry is 4.5^37^, lower than that of eGFP (pKa 6.0)^38^, mCherry is more resistant to quenching in acidic environments. Thus, autophagosome maturation can be monitored using this mCh-eGFP-LC3B marker; newly formed, immature autophagosomes are marked with both red and green fluorescence, while mature autolysosomes with acidified lumens emit only mCh fluorescence (Figure 2A)^36^.

**Figure 2.**
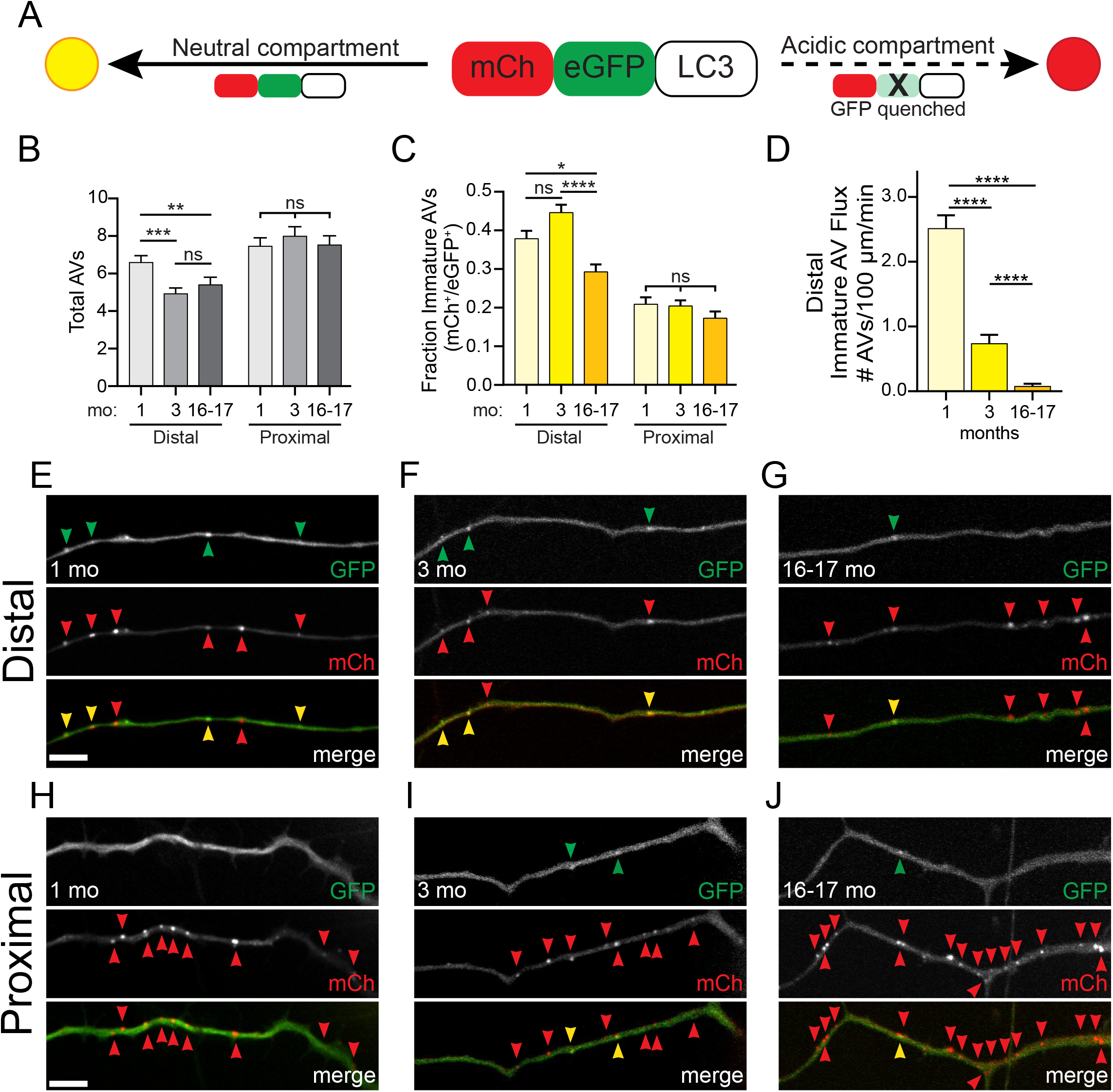
Autophagic vesicle maturation increases in the distal axon during aging. (A) Schematic of the tandem mCh-eGFP-LC3B probe in an immature autophagosome (top) and an acidified, mature autolysosome (bottom). (B) Quantification of the total number of autophagic vesicles present in the distal (left) and proximal (right) axon in DRG neurons from young, young adult, and aged mice (mean ± SEM; n≥ 80 neurons from three biological replicates), normalized to axon length and imaging duration. **p < 0.005; ***p < 0.0005; ns, not significant by Kruskal-Wallis test with Dunn’s multiple comparisons test. (C) Quantification of the fraction of GFP^+^/mCh^+^ autophagic vesicles in the distal (left) and proximal (right) axon in DRG neurons from young, young adult, and aged mice (mean ± SEM; n≥ 79 neurons from three biological replicates). *p < 0.05; ****p < 0.0001; ns, not significant by Kruskal-Wallis test with Dunn’s multiple comparisons test. (D) Quantification of the number of GFP^+^/mCh^+^ AVs detected per minute per 100 μm in the distal axon of DRG neurons from young, young adult, and aged mice (mean ± SEM; n≥ 79 neurons from three biological replicates). ****p < 0.0001 by Kruskal-Wallis test with Dunn’s multiple comparisons test. (E-G) Representative micrographs mCh-eGFP-LC3B in the distal axon of DRG neurons from young (E), young adult (F), and aged (G) mice. Top micrograph is the GFP channel; middle micrograph is the mCh channel; and bottom micrograph is the merge. In the merge panels, yellow arrowheads denote an immature, GFP^+^/mCh^+^ autophagosome and red arrowheads denote a mature, GFP quenched, mCh^+^ autolysosome. Retrograde is to the right. Scale bar, 5 μm. (H-J) Representative micrographs mCh-eGFP-LC3B in the proximal axon of DRG neurons from young (H), young adult (I), and aged (J) mice. Top micrograph is the GFP channel; middle micrograph is the mCh channel; and bottom micrograph is the merge. In the merge panels, yellow arrowheads denote an immature, GFP^+^/mCh^+^ autophagosome and red arrowheads denote a mature, GFP quenched, mCh^+^ autolysosome. Retrograde is to the right. Scale bar, 5 μm.

Using the mCh-eGFP-LC3B tandem probe transiently transfected into primary DRG neurons, we first determined the absolute number of AVs, independent of maturation state, in the distal and proximal axon of DRG neurons during aging. Interestingly, we observed a significant decrease in the number of AVs in the distal axon during development, but not during aging. We also detected no age-related changes in the total number of AVs in the proximal axon (Figure 2B). We next assessed the percentage of AVs that were positive for both mCh and eGFP in the distal and proximal axon. At all ages examined, we saw a significantly higher fraction of immature AVs, positive for both mCh and eGFP fluorescence, in the distal axon as compared to the proximal axon (Figure 2C, 2E-J). These data indicate that aging does not disrupt the overall pathway for maturation of axonal autophagosomes. However, in neurons from aged mice, we detected a decrease in the fraction of immature AVs in the distal axon, but no age-related changes in AV maturation in the proximal axon (Figure 2C).

We next used the transiently transfected tandem probe to ask whether we could replicate the age-related decrease in autophagosome transport flux we observed using DRG neurons from GFP-LC3B transgenic mice (Figure 1G). Indeed, we observed significant decreases in immature AV transport flux in the distal axon between DRG neurons from young, young adult and aged mice with the ectopically expressed mCh-eGFP-LC3B marker (Figure 2D). It is important to note that the GFP-LC3B probe was expressed stably at a low level in neurons isolated from B6.Cg-Tg(CAG-EGFP/LC3)53Nmi/NmiRbrc transgenic mice^15,16,27,39^, while the mCh-eGFP-LC3B tandem probe was transiently transfected into DRG neurons after isolation. Higher expression levels of the dual reporter construct relative to the GFP-LC3B probe likely drive the higher number of total AVs observed in these experiments. Collectively, these results explain our observations of AV transport flux using the GFP-LC3B probe (Figure 1G), showing a decrease in total AV transport flux in neurons from young to young adult mice with a subsequent decrease in the percentage of distal immature AVs in neurons from young adult to aged mice.

### Lysosomal Content of the Distal Axon does not Change with Age

We next investigated the molecular and cellular underpinnings of the decrease in the fraction of immature AVs in the distal axon during aging. The age-related change in AV maturation that we observed in the distal axon could be due to alterations in lysosomal content or in AV transport in the distal axon. To examine the first possibility, we assessed multiple parameters of lysosomes in the distal axon of neurons from three ages of mice. We first used LysoTracker (LysoT) Red to label acidic vesicles and CellMask Deep Red to label the plasma membrane (Figure 3A). We acquired z-stacks of the distal tip of axons and then calculated the number of LysoT-positive puncta (Figure 3B) and LysoT-positive area (Figure 3C), both normalized to the area of the distal axon. We observed no significant changes with age in either normalized number or area of LysoT+ puncta in the distal axon. Second, we used Cresyl Violet (CresylV, ^40^) as an alternative label of acidic vesicles, again in parallel with CellMask Deep Red labeling of the plasma membrane (Figure 3D). We observed no age-related alterations in normalized number of CresylV-positive puncta in the distal axon (Figure 3E). While we observed a decrease in the normalized area of CresylV-positive vesicles in the distal axon between neurons from young and young adult mice, we did not observe any changes in normalized area of CresylV-positive puncta during aging (Figure 3F). Furthermore, this observed decrease in normalized area does not explain the detected age-related increase in AV maturation in the distal axon. These data suggest there are no age-related increases in acidic lysosomes and late endosomes in the distal axon of DRG neurons.

**Figure 3.**
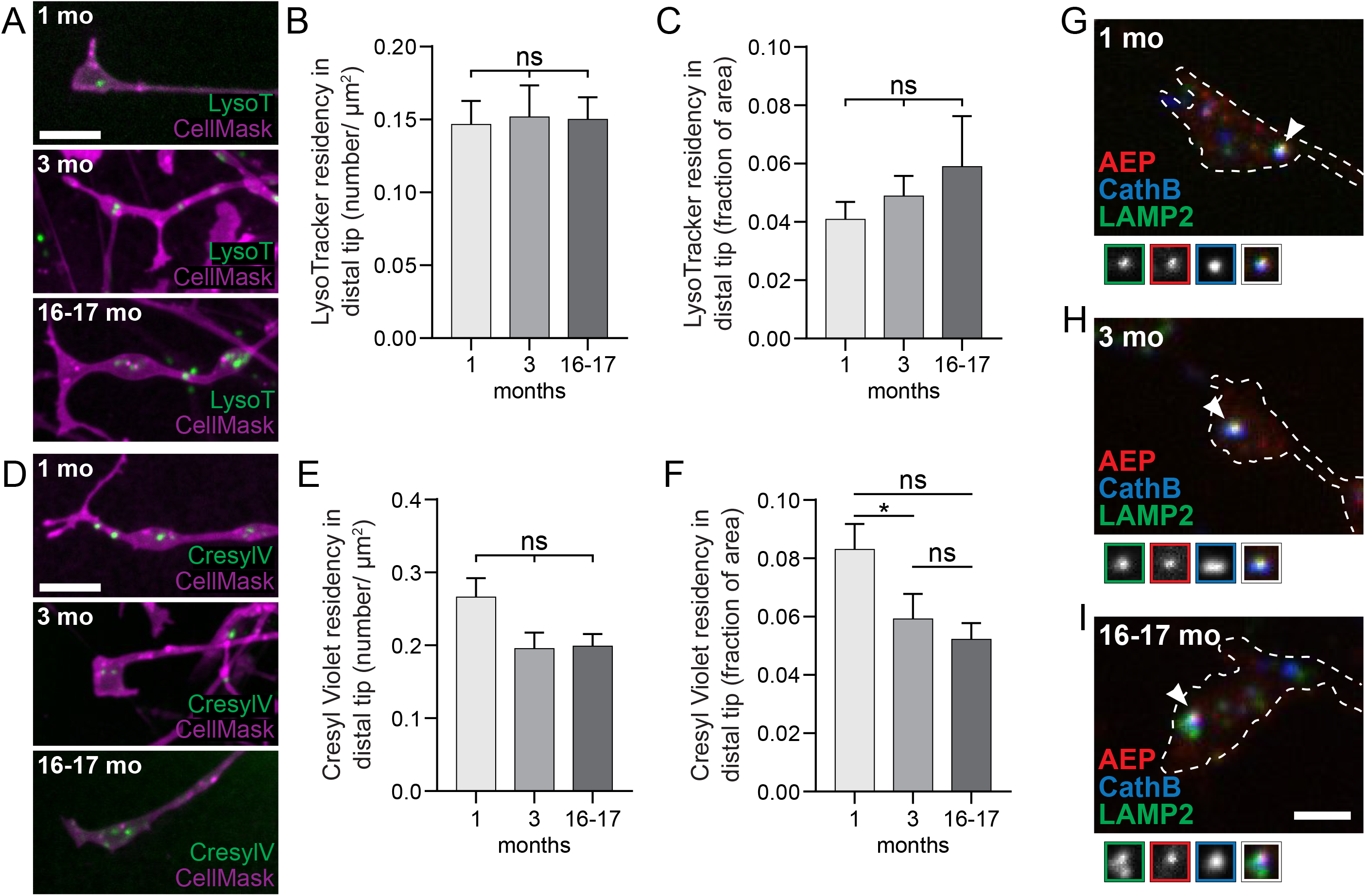
Lysosomal content in the distal axon does not change with age. (A) Representative maximal projection micrographs of LysoTracker Red (“LysoT”, green) and Cell Mask Deep Red (“Cell Mask”, magenta) from live cell imaging of the distal neurite of DRG neurons from young (top), young adult (middle), and aged (bottom) mice. Scale bar, 5 μm. (B-C) Quantification of the number of LysoTracker Red-positive puncta (B) and the area of LysoTracker Red-positive puncta (C), normalized to distal tip area in DRG neurons from young (light gray), young adult (gray), and aged (dark gray) mice (mean ± SEM; n≥ 83 neurons from three biological replicates ns, not significant by Kruskal-Wallis ANOVA test with Dunn’s multiple comparisons test. (D) Representative maximal projection micrographs of Cresyl Violet (“CresylV”, green) and Cell Mask Deep Red (“Cell Mask”, magenta) from live cell imaging of the distal neurite of DRG neurons from young (top), young adult (middle), and aged (bottom) mice. Scale bar, 5 μm. (E-F) Quantification of the number of Cresyl Violet-positive puncta (E) and the area of Cresyl Violet-positive puncta (F), normalized to distal tip area in DRG neurons from young (light gray), young adult (gray), and aged (dark gray) mice (mean ± SEM; n≥ 87 neurons from three biological replicates). *p < 0.05; ns, not significant by Kruskal-Wallis ANOVA test with Dunn’s multiple comparisons test. (G-I) Representative maximal projection micrographs of the distal neurites of fixed DRG neurons from young (G), young adult (H) and aged (I) mice. White arrowheads indicate colocalization of anti-AEP (red), anti-Cathepsin B (blue, “CathB”), and anti-LAMP2 (green). Borders of magnifications of indicated puncta denote channel (left 3 grayscale boxes) or colocalization state in merge (right box). Scale bar, 2 μm.

Degradatively competent lysosomes are present in the distal axon of primary hippocampal, cortical, and DRG neurons^41–44^. Thus, we asked whether we could detect any age-related changes in the presence of lysosomal enzymes in the distal axon. We used multi-color immunocytochemistry to examine the localization of endogenous lysosomal enzymes lysosomal associated protein 2 (LAMP2), Cathepsin B, and Asparagine Endopeptidase (AEP) in DRG neurons from young (Figure 3G), young adult (Figure 3H) and aged (Figure 3I) mice. We observed colocalization of all three lysosomal enzymes in the distal axon of neurons from all three ages of mice, indicating that lysosomes containing degradative enzymes are present in the distal axon during development and aging. Next, we examined the endogenous levels of lysosomal proteins by immunoblot of whole brain lysates. We did not detect age-related changes in the levels of lysosomal proteins lysosomal associated protein 1 (LAMP1), lysosomal integral membrane protein 2 (LIMP2), Cathepsin D, and lysosomal H+ V-type ATPase subunit A (ATP6V1A) in whole mouse brain lysate (Figure S1). We did detect modest increases in the levels of Cathepsin L and lysosomal H+ V-type ATPase subunit B (ATP6V1B) in whole brain lysates from aged mice (Figure S1). However, increases in individual lysosomal enzymes or subunits of the H+ ATPase likely cannot fully account for the age-associated alterations we observed in AV maturation. Taken together, our data suggest that neither lysosomal residency nor degradative capacity changes in the distal axon with age, implying that the age-related changes with observed in autophagosome maturation and transport flux are not due to age-related alternations in lysosomal capacity in the distal axon.

### Many Parameters of AV Transport in the Distal Axon do not Change with Age

We next asked whether AV transport changed with age. To assess AV transport, we used multicolor, live-cell microscopy to acquire 3-minute time-lapse videos of DRG neurons transiently expressing the tandem mCh-eGFP-LC3B probe. We chose the tandem probe so that we could analyze axonal transport of both immature, mCh^+^/eGFP^+^ autophagosomes and mature, mCh^+^/eGFP^-^ autolysosomes. By analyzing the kymographs (Figure 4A) generated from the time-lapse videos, we calculated several parameters of axonal AV transport. Each AV was categorized based on its net displacement during imaging: “anterograde” if the AV net displacement was greater than 10 μm toward the axonal tip, “bidirectional/stationary” if the AV net displacement was less than 10 μm, and “retrograde” if the AV net displacement was greater than 10 μm toward the cell soma (Figure S2).

**Figure 4.**
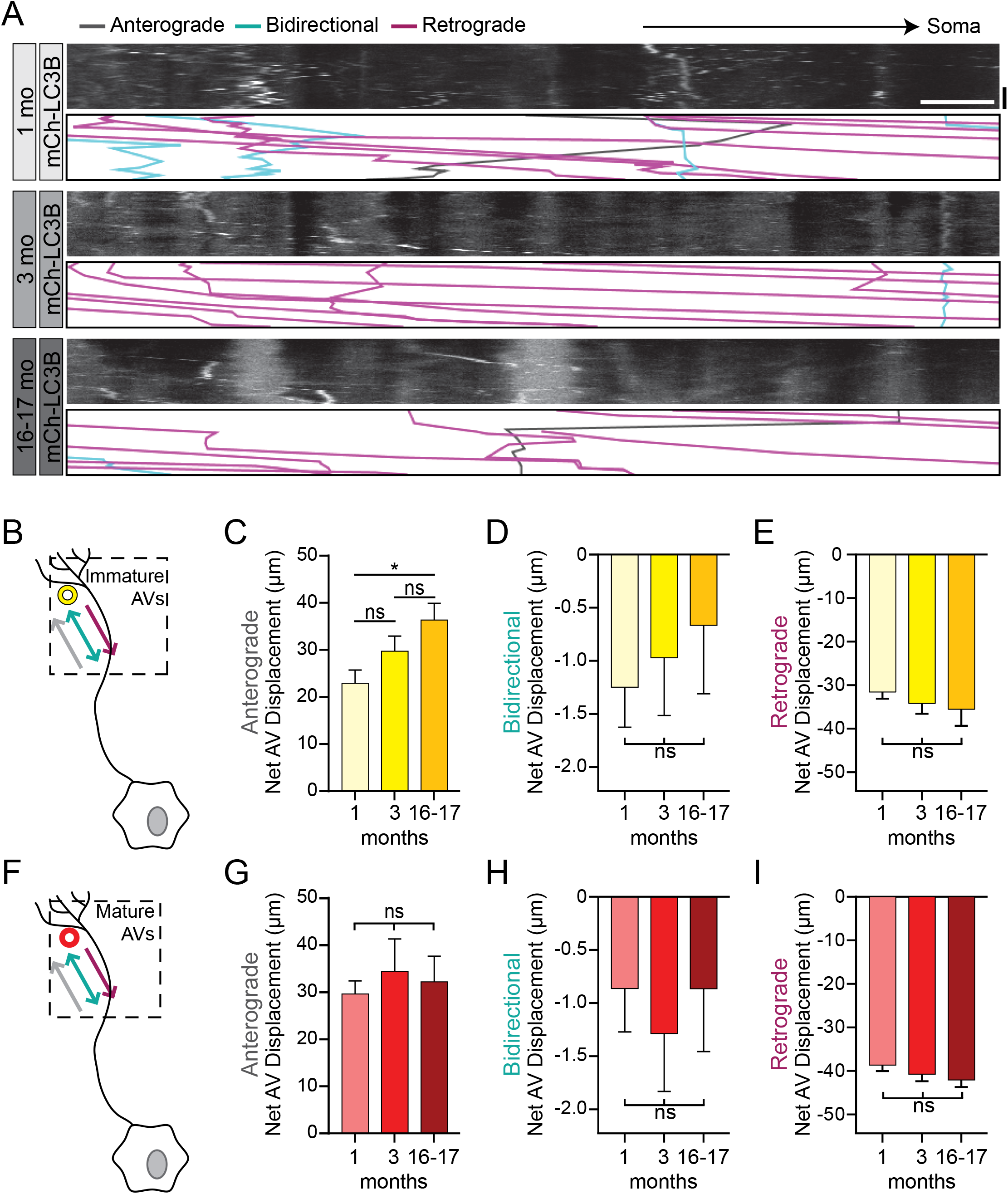
Transport dynamics of immature and mature AVs do not considerably change with age in the distal neurite. (A) Representative kymographs of the mCh channel of the distal neurite of DRG neurons ectopically expressing mCh-eGFP-LC3B from young (light gray), young adult (gray), and aged (dark gray) mice. Annotations of individual AV tracks are below the grayscale kymographs. Gray tracks represent puncta that moved in a net anterograde direction; cyan tracks represent puncta that moved in a net bidirectional direction; and magenta tracks represent puncta that moved in a net retrograde direction. Retrograde is to the right. Horizontal scale bar, 5 μm; vertical scale bar, 1 min. (B) Cartoon of DRG neuron depicting time-lapse imaging of immature AVs in the distal neurite. (C-E) Quantification of the net displacement of immature autophagic vesicles that moved in a net anterograde (C), net bidirectional or stationary (D) or net retrograde (E) direction in the distal neurite of DRG neurons from young (1 mo, light yellow), young adult (3 mo, yellow), and aged (16-17 mo, dark yellow) mice (mean ± SEM; n≥ 18 puncta from three biological replicates). *p < 0.05; ns, not significant by Kruskal-Wallis test with Dunn’s multiple comparisons test. (F) Cartoon of DRG neuron depicting time-lapse imaging of mature AVs in the distal neurite. (G-I) Quantification of the net displacement of mature autophagic vesicles that moved in a net anterograde (G), net bidirectional or stationary (H) or net retrograde (I) direction in the distal neurite of DRG neurons from young (light red), young adult (red), and aged (dark red) mice (mean ± SEM; n≥ 15 puncta from three biological replicates). ns, not significant by Kruskal-Wallis test with Dunn’s multiple comparisons test.

We first analyzed the transport dynamics of immature, mCh^+^/eGFP^+^ autophagosomes in the distal axon. While there were no age-related changes in net displacement for bidirectional or retrograde AVs, we did note a significant increase in net displacement of anterograde AVs when comparing neurons from young and aged mice (Figure 4B-E). We did not observe age-related changes in average speed (Figure S3A-D) or pause time fraction (the time spent not moving/total time of AV run) across conditions (Figure S4A-D). Additionally, we did not observe significant age-related changes in total run length, pause duration, or pause number for anterograde, bidirectional or retrograde immature AVs (data not shown). These data indicate that most transport parameters for immature autophagosomes in the distal axon are not significantly altered with age.

We next analyzed the transport dynamics of mature, mCh^+^/eGFP^-^ autolysosomes in the distal axon. We did not detect any age-related changes in AV net displacement for anterograde, bidirectional, or retrograde AVs (Figure 4F-I). Similarly, we did not observe consistent changes in average speed with age, although we did detect a slight decrease in the average speed of bidirectional AVs in neurons from aged mice compared to young mice and an increase in average speed in retrograde AVs in neurons from young adult compared to young mice (Figure S3E-H). While we did observe an increase in the AV pause time fraction in bidirectional AVs in the distal axon in neurons from adult mice compared to young mice, we detected no age-related changes in AV pause time fraction in anterograde or retrograde AVs in the distal axon (Figure S4E-H). Additionally, we did not observe significant age-related changes in total run length, pause duration, or pause number for anterograde or retrograde AVs (data not shown). Collectively, our data indicate that AV transport parameters for both immature and mature AVs are generally unchanged in the distal axon with age. These data further suggest that the overall capability of axonal microtubule motors, kinesins and cytoplasmic dynein, is maintained during development and aging.

### Transport of autophagic vesicles becomes more efficient with age in the distal axon

Interestingly, we did identify one AV transport parameter that broadly changed with age in the distal axon. When we analyzed the number of times an AV switched directions during the imaging video (“switch count”), we observed a significant decrease with age from young to young adult mice that was subsequently preserved during aging. For immature AVs, we detected 51% and 52% decreases in AV switch count in the distal axon of neurons between young and young adult mice for anterograde and retrograde AVs, respectively. Furthermore, this decrease was sustained during aging (Figure 5A-D). Similarly, for mature AVs, we noted 55% and 57% decreases in AV switch count between neurons from young and young adult mice for bidirectional and retrograde AVs, respectively (Figure 5F-I). Likewise, this decrease was subsequently retained during aging (Figure 5F-I). Both kinesins and cytoplasmic dynein are simultaneously bound to axonal AVs^15,41,45^. Motor adaptor proteins are also present on neuronal AVs and regulate the balance between anterograde and retrograde movement by modulating the activation state of kinesin and dynein motors^20,41^. Thus, our data suggest that the regulation of the molecular motors bound to axonal AVs may change between neurons from 1-month and 3-month mice to enable fewer directional switches during AV axonal transport.

**Figure 5.**
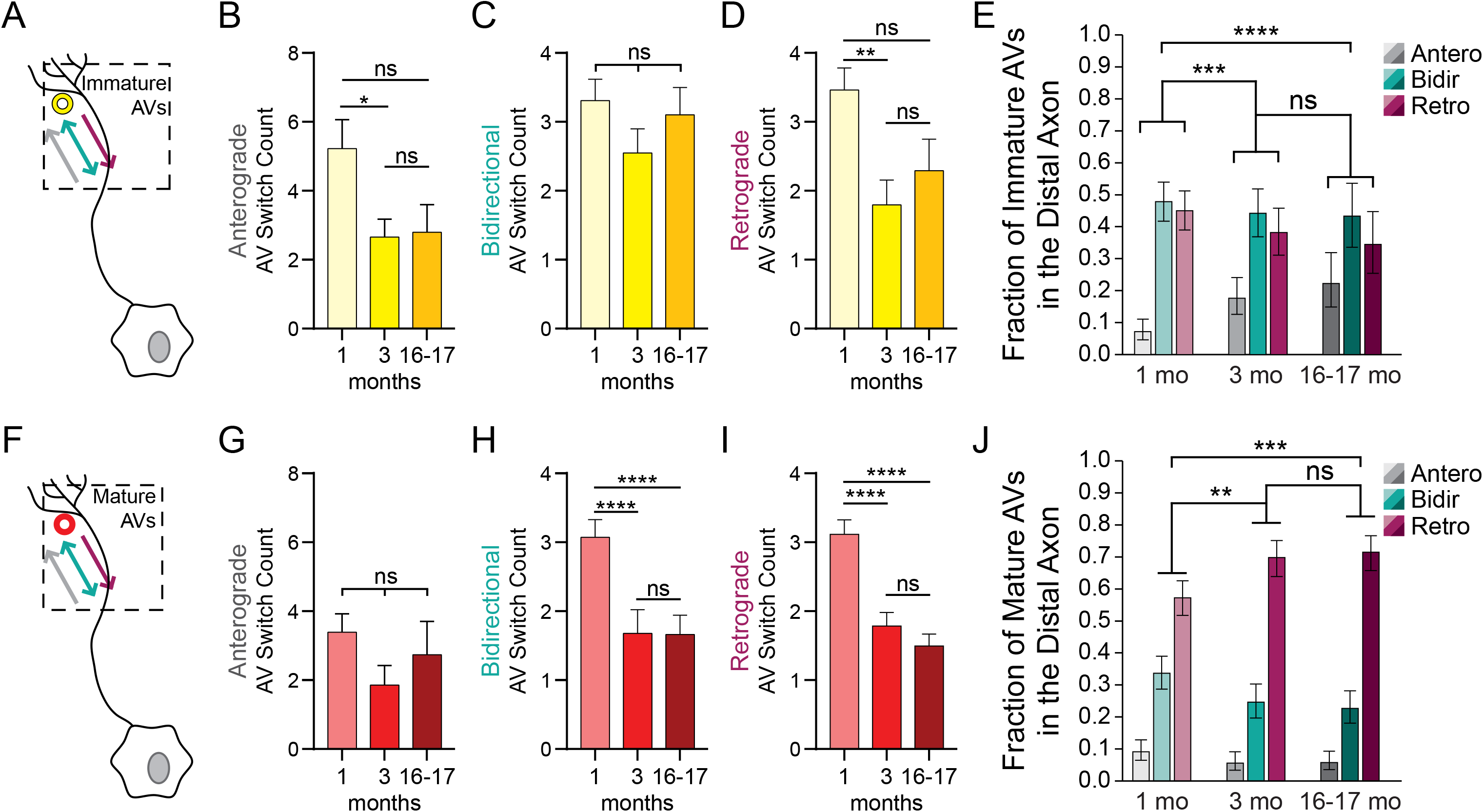
Transport of immature and mature autophagic vesicles in the distal axon becomes more efficient during development. (A) Cartoon of DRG neuron depicting time-lapse imaging of immature AVs in the distal neurite. (B-D) Quantification of the switch count of immature autophagic vesicles that moved in a net anterograde (B), net bidirectional or stationary (C) or net retrograde (D) direction in the distal neurite of DRG neurons from young (1 mo, light yellow), young adult (3 mo, yellow), and aged (16-17 mo, dark yellow) mice (mean ± SEM; n≥ 18 puncta from three biological replicates). **p < 0.005; *p < 0.05; ns, not significant by Kruskal-Wallis test with Dunn’s multiple comparisons test. (E) Quantification of the fraction of immature AVs in the distal axon that moved in a net anterograde (gray), bidirectional/stationary (cyan), or retrograde (magenta) direction from young (1 mo), young adult (3 mo), and aged (16-17 mo) mice. ***p=0.0016 by Chi-square test (mean ± 95% confidence intervals; n≥ 18 puncta from three biological replicates). ***p < 0.005; ****p < 0.0001; ns, not significant by Fisher’s exact test. (F) Cartoon of DRG neuron depicting timelapse imaging of mature AVs in the distal neurite. (G-I) Quantification of the net displacement of mature autophagic vesicles that moved in a net anterograde (G), net bidirectional or stationary (H) or net retrograde (I) direction in the distal neurite of DRG neurons from young (light red), young adult (red), and aged (dark red) mice (mean ± SEM; n≥ 15 puncta from three biological replicates). ****p < 0.0001; ns, not significant by Kruskal-Wallis test with Dunn’s multiple comparisons test. (J) Quantification of the fraction of mature AVs in the distal axon that moved in a net anterograde (gray), bidirectional/stationary (cyan), or retrograde (magenta) direction from young (1 mo), young adult (3 mo), and aged (16-17 mo) mice. ***p=0.0028 by Chi-square test (mean ± 95% confidence intervals; n≥ 14 puncta from three biological replicates). **p < 0.01; ***p < 0.005; ns, not significant by Fisher’s exact test.

We also examined the fraction of AVs that moved in a net anterograde, bidirectional, or retrograde direction. For immature AVs in the distal axon, independent of age, the plurality of AVs moved bidirectionally, with a slightly lower fraction moving in a net retrograde direction (Figure 5E). Of interest, the fraction of immature AVs that moved in a net anterograde direction increased between neurons from young and young adult mice and was maintained during aging (Figure 5E). This increase in the anterograde fraction was accompanied by a concomitant slight decrease in the retrograde fraction (Figure 5E). These data suggest that there are modifications during development in the distal axon that result in immature AVs being held in the distal axon and these modulations are preserved during aging.

Similarly, we assessed the net direction of movement of mature AVs in the distal axon. Independent of age, the majority of mature AVs in the distal axon moved in a net Retrograde direction (Figure 5J). In contrast to the immature AVs in the distal axon, we observed an increase in the fraction of mature AVs that moved in a net retrograde direction between neurons from 1-month and 3-months which was retained during aging (Figure 5J). This age-related increase in the retrograde fraction was balanced by a concomitant decrease in the fraction of mature AVs that moved bidirectionally. These data suggest that age-related modifications in motor adaptor proteins also influence the direction of net movement of mature AVs in the distal axon. Taken together, our data suggest that a sorting mechanism for AVs is refined in the distal axon during development that holds immature AVs in the distal axon while simultaneously enabling robust retrograde transport of mature AVs out of the distal compartment. Importantly, the efficacy of this distal sorting mechanism is sustained during aging.

### Retrograde Transport of Mature Autolysosomes in the Proximal Axon Becomes More Efficient with Age

We next assessed AV transport in the proximal axon. Approximately 80% of AVs in the proximal are mature, mCh^+^/eGFP^-^ autolysosomes (Figure 2C). Independent of age, in both the distal and proximal axon, the majority of mature AVs are transported in a net retrograde direction (Figure S2). We calculated the transport parameters for each category of AVs (immature and mature) in the proximal axon: anterograde, bidirectional, and retrograde. However, due to the small numbers of immature AVs moving in any direction and mature AVs moving net anterogradely or bidirectionally (Figure S2), we focused on the mature AVs moving retrogradely in the proximal axon (Figure 6A-B). Comparing neurons from young and young adult mice, we observed an increase in total run length (Figure 6C), net displacement (Figure 6D), and average speed (Figure 6E). Additionally, we observed decreases in pause time fraction (Figure 6F) and switch count (Figure 6G) between neurons from young and young adult mice. These changes were generally maintained during aging (Figure 6C-G), with only average speed and switch count differing significantly between neurons from young adult and aged mice (Figure 6E, 6G). These data diverge from our observations of few age-related changes in AV transport parameters in the distal axon and suggest that age-related changes in the transport dynamics of mature, retrogradely moving AVs occur primarily in the proximal axon. However, our findings in both the distal and proximal axon are congruent with an axonal sorting mechanism for AVs that becomes more efficient during development and is sustained during aging.

**Figure 6.**
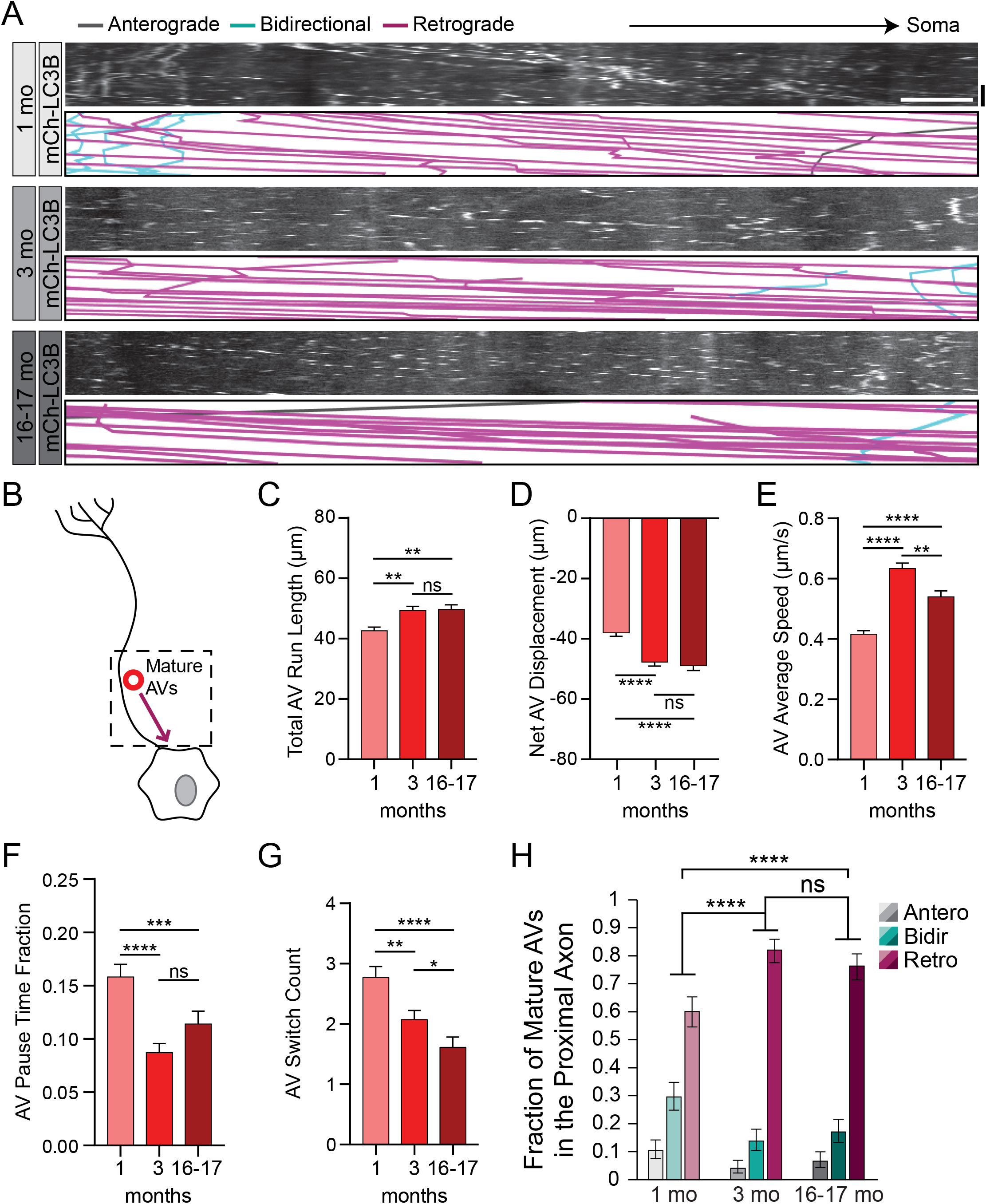
Retrograde transport of mature autophagic vesicles in the proximal axon becomes more efficient during development. (A) Representative kymographs of the mCh channel of the proximal neurite of DRG neurons ectopically expressing mCh-eGFP-LC3B. Annotations of individual AV tracks are below the grayscale kymographs. Retrograde is to the right. Horizontal scale bar, 5 μm; vertical scale bar, 1 min. (B) Cartoon of DRG neuron depicting time-lapse imaging of immature AVs in the proximal neurite. (C-G) Quantification of the total run length (C), net displacement (D), average speed (E), pause time fraction (F), and switch count (G) for mature AVs that had a net retrograde displacement in the proximal axon of DRG neurons (mean ± SEM; n≥ 242 puncta from three biological replicates). ****p < 0.0001; ***p < 0.005; **p < 0.01; *p < 0.05; ns, not significant by Kruskal-Wallis test with Dunn’s multiple comparisons test. (H) Quantification of the fraction of mature AVs in the proximal axon that moved in a net anterograde (gray), bidirectional/stationary (cyan), or retrograde (magenta) direction from young (1 mo), young adult (3 mo), and aged (16-17 mo) mice. ****p < 0.0001 by Chi-square test (mean ± 95% confidence intervals; n≥ 13 puncta from three biological replicates). ****p < 0.0001; ns, not significant by Fisher’s exact test.

Additionally, we analyzed the net direction of movement of mature AVs in the proximal axon. Similar to the distal axon, independent of age, the majority of mature AVs were transported in a net retrograde direction in the proximal axon (Figure 6H). Also similar to the distal axon, we observed an increase in the fraction of mature AVs that moved in a net retrograde direction with a simultaneous decrease in the fraction that moved bidirectionally between neurons from young and young adult mice. These alterations were retained during aging, as we detected no changes between neurons from young adult and aged mice (Figure 6H). Collectively, these data in the proximal axon mirror our data from the distal axon, in which a sorting mechanism is consolidated during development and preserved during aging to efficiently move mature autolysosomes through the proximal axon into the cell soma to enable effective recycling of neuronal AV contents into new proteins and organelles.

Together, these data extend the current model for axonal transport of autophagic vesicles^41^ to focus on changes during development and aging. During development, axonal compartments establish sorting mechanisms for immature and mature autophagic vesicles, restricting immature autophagosomes distally and releasing mature autolysosomes to move proximally (Figure 7). This sorting capability is retained in neurons from aged mice, suggesting that aged neurons would have the capacity to effectively process AVs if autophagosome biogenesis was restored.

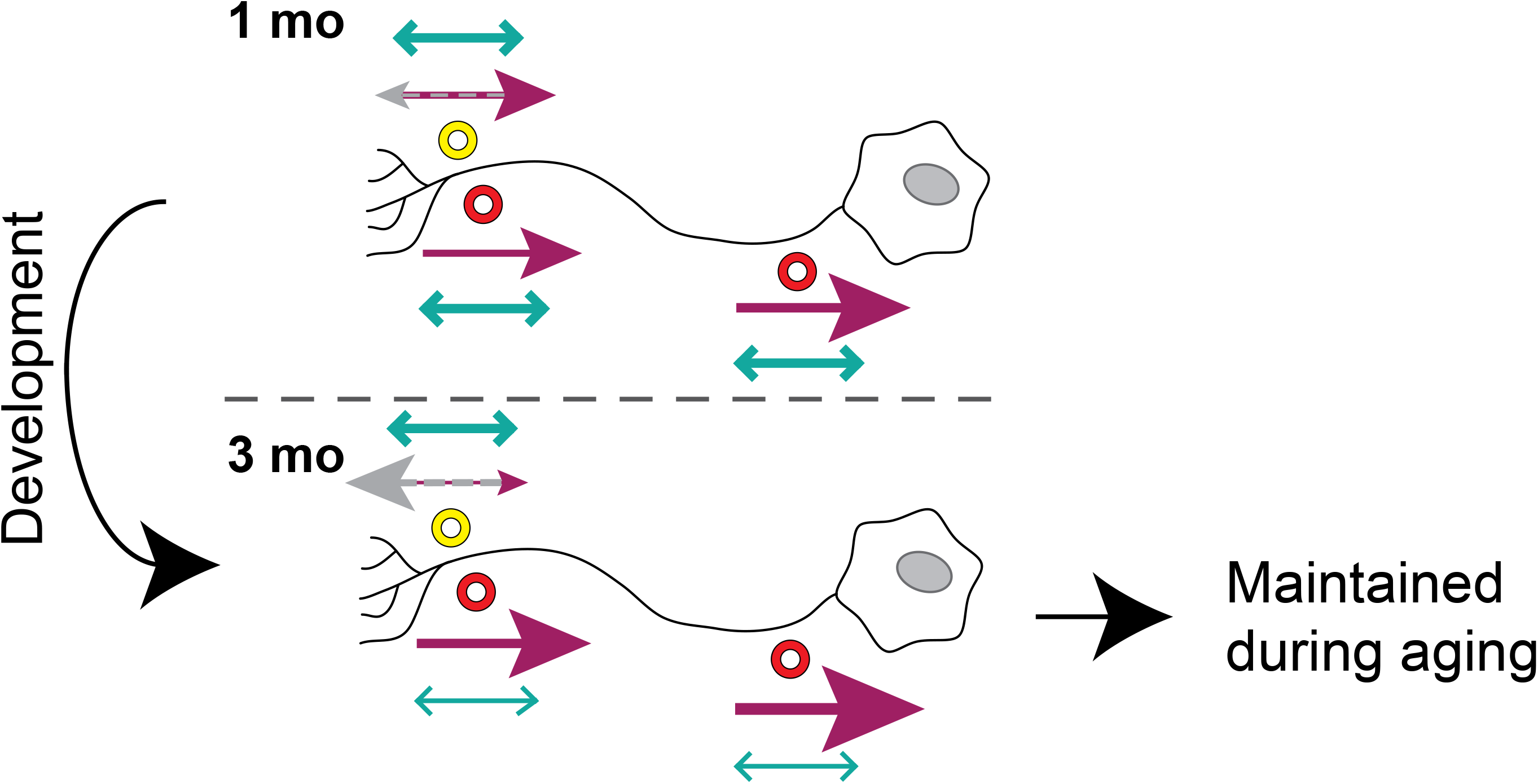

## DISCUSSION

We have examined the dynamics of axonal autophagy during aging and shown that while the rate of autophagosome biogenesis decreases with age, transport of autophagosomes and autolysosomes is not globally impaired during aging. Conversely, we have demonstrated that transport of mature AVs in the proximal axon actually becomes more efficient during development and that this efficiency is maintained during early aging. Interestingly, while we observed an age-related decrease in AV transport flux in the mid-axon with the GFP-LC3B autophagosomal reporter, we determined that this decrease was likely due to a significant decrease in the rate of autophagosome biogenesis combined with an increase in the maturation of AVs in the distal axon during aging. Ultimately, we demonstrated that DRG neurons from aged mice do not have a significant reduction in AV maturation nor a diminished capacity to transport AVs in the distal or proximal axon, suggesting that the later stages of autophagy remain robust in neurons during aging.

Ectopic expression of autophagy pathway components has been shown to affect the rate of autophagosome biogenesis. Overexpression of LC3B can enhance autophagosome biogenesis^46–48^, consistent with the higher flux measurements observed in experiments overexpressing the tandem reporter probe (Figure 2D) compared to data collected using the transgenic GFP-LC3B reporter (Figure 1G). We found that by ectopically expressing the tandem mCh-eGFP-LC3B AV reporter, we eliminated the variable of age-related changes in AV number per minute in the axon. With ectopic expression, we observed no age-related changes in AV number in the proximal axon and an age-related decrease of only one AV per minute in the distal axon (Figure 2B). Thus, in our experiments examining AV transport dynamics, the neurons from young, young adult, and aged mice were transporting approximately the same number of AVs in the distal and proximal axon, further suggesting that neurons preserve their capacity to transport AVs efficiently during aging.

Given our data indicating that autophagosomes mature into autolysosomes more rapidly in the distal axon in neurons from aged mice relative to neurons from young and young adult mice (Figure 2C), we hypothesized that this age-related increase in autophagosome maturation could be due to an age-induced increase in endolysosomal content in the distal axon or to changes in AV transport dynamics. Lysosomes and late endosomes fuse with autophagosomes to generate autolysosomes, which eventually degrade and recycle engulfed AV cargo. Since the endolysosomal system directly regulates the maturity of AVs, we sought to identify any age-related alterations in lysosomal content in the distal axon. Using acid-sensitive dyes and interrogating endogenous endolysosomal proteins, we did not detect age-related changes in lysosomal content or the presence of degradative enzymes in the distal axon. Importantly, we did not examine the degradative capacity of the acidified endolysosomes in the distal axon. Others have suggested that there is a gradient of degradative capability of endolysosomes in the axon, with more degradatively competent lysosomes positioned in the proximal axon^49^, although lysosomes containing degradative capability have been observed in the distal axon of primary neurons^42–44^. Since our initial results indicated age-related changes to AV maturation only in the distal axon, we did not investigate whether aging modulates the gradient of endolysosomal degradative capacity in the proximal axon of DRG neurons. Defects in the endolysosomal system have been implicated in age-related neurodegenerative diseases^50,51^. It is possible that our results suggesting that distal AVs acidify faster in neurons from aged mice could contribute to age-related defects in the endolysosomal pathway. However, acidification of endolysosomes is required for the activation of their degradative machinery; thus, we hypothesize that this larger fraction of acidified AVs indicates that aging alone does not inhibit progression through the later stages of autophagy in neurons. In the future, it will be important to assess how age modulates the degradative capacity of endolysosomes in distinct axonal regions and in models of NDDs.

While we found no evidence for alterations in endolyosomal presence in the distal axon (Figure 3), we did observe changes in specific parameters of AV transport in the distal axon. Despite the majority of AV transport parameters not changing with age in the distal axon for both immature and mature AVs (Figure 4), we did observe age-related decreases in the number of times AVs switched directions in the distal axon (Figure 5). Additionally, we found that the fraction of immature autophagosomes moving anterograde increased with age, while the fraction of mature autolysosomes moving retrograde increased with age. However, these changes occurred during development, observed by comparing neurons from young (1 mo) and young adult (3 mo) mice. Conversely, we observed an increase in distal AV maturation in neurons from aged mice (Figure 2C). Since the ectopic expression of the mCh-eGFP-LC3B tandem marker normalized the number of distal autophagosomes between neurons from young adult and aged mice, age-related changes in distal AV maturation are not the result of age-related changes in the proportion of lysosomes to autophagosomes in the distal axon. Therefore, our data suggest that there are other mechanisms that are modulated during aging that alter AV maturation state. One possible mechanism is that the fusion of autophagosomes to lysosomes becomes more efficient during aging, yielding a higher fraction of mature AVs in the distal axon in neurons from aged mice. It will be interesting to examine age-related changes in autophagosome-endolysosome fusion in future studies.

Interestingly, while we observed only AV switch count changing substantially with age in the distal axon (Figure 5), several AV transport parameters changed during development in the proximal axon (Figure 6). Together, our data from the distal and proximal axon suggest that the properties of the microtubule motors cytoplasmic dynein and kinesins are not altered with age, but instead that motor adapters are modulated during development and aging. Motor adapters not only link vesicles, including AVs, to the motor complexes, but also regulate the activation state of the cytoplasmic dynein and kinesin motors that are simultaneously bound to these organelles^20,52^. In embryonic primary neurons, different motor adapters predominate in different axonal compartments, which could explain how aging broadly affects AV transport parameters for mature AVs in the proximal axon while only specifically altering AV switch count in the distal axon^41^. Remarkably, we observed that AV sorting in the distal axon becomes more efficient during development, with a higher fraction of immature autophagosomes moving anterograde and a higher fraction of mature autolysosomes moving retrograde in neurons from young adult mice compared to neurons from young mice. This increased efficiency is retained during early aging and suggests a robust sorting mechanism for the transport of immature and mature AVs. Since motor adapters participate in cargo selection for microtubule motors,^52^ motor adapters are particularly attractive candidates for this sorting mechanism. While AV-associated motor adapters have recently been studied in the context of axonal transport in cortical and hippocampal embryonic neurons^41^, it will be important to examine the spatial and temporal dynamics of AV-relevant motor adapters in neurons during development and aging.

One compelling motor adaptor that might regulate this sorting mechanism is JNK-interacting protein 4 (JIP4). Overexpression of JIP4 in murine cortical neurons disrupted the retrograde transport of AVs, resulting in a higher proportion of AVs moving bidirectionally or anterogradely^53^. JIP4 can bind to RAB proteins that are phosphorylated by leucine-rich repeat kinase 2 (LRRK2)^54,55^, and JIP4 can recruit and activate kinesin^56^. Collectively, this suggests a model in which LRRK2 could recruit JIP4 to immature AVs, activating the anterograde motor kinesin and restricting those autophagosomes to the distal axon. Interestingly, mutations in LRRK2 are the most prevalent genetic cause of PD^57^, underscoring the role of neuronal autophagy in neurodegenerative disease. Another compelling motor adaptor is the structurally unrelated JIP1^58,59^. Phosphorylation of JIP1 S421 was shown to regulate autophagosome transport in DRG neurons by coordinating the activity of opposing kinesin and dynein motors; JIP1 is required for autophagosomal exit from the distal axon and the switch to processive retrograde transport in the mid axon^20^. Consistent with these findings, JIP1 preferentially associates with immature autophagosomes in primary rat hippocampal neurons^41^. Thus, the phosphorylation state of JIP1 on immature autophagosomes could contribute to the axonal autophagosomal sorting mechanism, particularly in the distal axon.

Misregulation of autophagy has been implicated in many age-related neurodegenerative diseases, with multiple stages of autophagy being affected in a specific disease and multiple diseases influencing a given stage of autophagy^4–6^. The most relevant shared risk factor for age-related neurodegenerative diseases is age^28^. Our data suggest that while autophagosome biogenesis dramatically decreases during aging (Figure 1), later stages of autophagy, including autophagosomal maturation, do not become less effective with age. Conversely, AV transport becomes more efficient during development and is maintained during aging, while autophagosome maturation in the distal axon occurs more rapidly in neurons from aged mice. Thus, our results imply that if we can restore the rate of autophagosome biogenesis in aged neurons, those neurons will be able to process and transport autophagosomes to eventually recycle the contents in the cell soma. For neurodegenerative diseases with known defects in autophagosomal transport and/or maturation, it will be important to address the synergistic effects of both age and disease.

Autophagy disfunction has also been associated with neuropathic pain^60–62^. Further, the incidence of neuropathic pain is correlated with age and is connected with certain neurodegenerative diseases. Studies in DRG neurons in models of neuropathic pain have observed increased autophagy after nerve injury^60,63–66^. Furthermore, increasing autophagy via rapamycin treatment alleviates neuropathic pain in mice^64^. Thus, our results demonstrating that autophagosome biogenesis decreases with age (Figure 1 & ^27^) suggest that the age-related decrease in autophagosome formation in DRG neurons could directly contribute to increased neuropathic pain during aging. Fortunately, our data also suggest that if autophagosome biogenesis can be augmented in aged neurons, the neurons will retain their ability to transport the AVs back to the cell soma to allow for degradation and recycling of cargo. Our previous work indicates that increasing the proportion of dephosphorylated WIPI2B in neurons from aged mice restores the age-related decrease in neuronal autophagosome biogenesis^27^, identifying a potential therapeutic avenue for treating age-related neuropathic pain. Ultimately, our data suggest that while axonal autophagosome biogenesis decreases substantially during aging, maturation and transport of AVs do not decline during aging, simplifying the therapeutic landscape to ameliorate autophagy-related diseases and conditions in neurons.

## ACKNOWLEDGEMENTS

HT was supported by R00 NS109286. ELFH was supported by R01 NS060698 and R37 NS060698. AKHS was supported by R00 NS109286, K99 NS109286 and R37 NS060698.

## METHODS

### Reagents

GFP-LC3B transgenic mice (strain: B6.Cg-Tg(CAG-EGFP/LC3)53Nmi/NmiRbrc) were generated by N. Mizushima (Tokyo Medical and Dental University, Tokyo, Japan; ^39^) and obtained from RIKEN BioResource Center in Japan. These mice were bred with C57BL/6J mice obtained from The Jackson Laboratory. Hemizygous and wild type littermates were used in experiments. Constructs used include: mCherry-ATG13 (subcloned from Addgene 22875), mCherry-GFP-LC3B (gift from T. Johansen, University of Tromsø, Tromsø, Norway; ^67^), and Halo-ATG14L (subcloned from Addgene 21635).

### Primary Neuron Culture

Mice were euthanized prior to dissection. All animal protocols were approved by the Institutional Animal Care and Use Committee at the University of Pennsylvania. DRG neurons were isolated as previously described^68^ from mice of either sex in these postnatal ranges: P21-28 (1 mo, young), P90-120 (3 mo, young adult), or P480-540 (16-17 mo, aged). DRG neurons were plated on glass-bottomed dishes (MatTek Corporation) and cultured in F-12 Ham’s media (Invitrogen) with 10% heat-inactivated fetal bovine serum, 100 U/mL penicillin, and 100 μg/mL streptomycin. DRG neurons were imaged or fixed after being maintained for 2 days at 37°C in a 5% CO_2_ incubator.

Prior to plating, neurons were transfected with a maximum of 0.6 μg total plasmid DNA using a Nucleofector (Lonza) and following the manufacturer’s instructions. For Halo-ATG14 transfected neurons, DRG neurons were incubated with 100nM of JF646-Halo ligand (from Luke Levis, Janelia Farms, HHMI) for at least 30 min at 37°C in a 5% CO2 incubator. After incubation, neurons were washed three times with complete equilibrated F-12 media, with the final wash remaining on the neurons for at least 15 min at 37°C in a 5% CO_2_ incubator.

### Immunofluorescence

DRGs were isolated, plated, and cultured as described above. At DIV2, DRG neurons were fixed in pre-warmed 50% Bouin’s solution with 4% sucrose in PBS for 30 minutes at room temperature. Neurons were washed three times with 1X PBS, then incubated in Cell Block (1X PBS with 1% BSA and 5% normal goat serum) with 0.4% Saponin (Sigma) for one hour at room temperature. DRG neurons were then incubated in Cell Block with 0.1% Saponin containing primary antibodies for one hour at room temperature. After three five-minute washes in 1X PBS, neurons were incubated in Cell Block containing secondary antibodies for one hour at room temperature. DRG neurons were washed three additional times in 1X PBS and then once with ddH2O. Neurons were then mounted in Prolong Gold, cured overnight in the dark at room temperature, and assessed by spinning disk confocal microscopy. See Key Resources Table for antibodies used.

### Live-cell imaging and image analysis

Microscopy was performed in low fluorescence nutrient media (Hibernate A, BrainBits) supplemented with 2% B27 and 2 mM GlutaMAX. Confocal images were captured with a spinning-disk confocal (UltraVIEW VoX; PerkinElmer) microscope (Eclipse Ti; Nikon) with an Apochromat 100x, 1.49 NA oil immersion objective (Nikon) at 37°C in an environmental chamber. The Perfect Focus System was used to maintain Z position during time-lapse acquisition. Digital micrographs were acquired with an EM charge-coupled device camera (C9100; Hammamatsu Photonics) using Volocity software (PerkinElmer).

To capture autophagosome biogenesis, time-lapse videos were acquired for 10 min with a frame every 3 sec. To capture autophagosome transport, time-lapse videos were acquired for 3 min with a frame every 3 sec. For each imaged neuron, a time-lapse video was acquired in the distal axon and then a subsequent time-lapse video was acquired in the proximal axon. Multiple channels were acquired consecutively, with the green (488 nm) channel captured first, followed by red (561 nm), and far-red (640 nm). DRG neurons were selected for imaging based on morphological criteria and low expression of transfected constructs. To minimize artifacts from overexpression, neurons within a narrow range of low fluorescence intensity were chosen for imaging, ensuring the analyzed neurons expressed low levels of the ectopic tagged proteins.

All image analysis was performed on raw data. Images were prepared in FIJI^69^; contrast and brightness were adjusted equally to all images within a series. Time-lapse micrographs were analyzed with FIJI^69^. To quantify AV biogenesis, GFP-LC3B puncta were tracked manually using FIJI. An AV biogenesis event was defined as the de novo appearance of a GFP-LC3B punctum based on changes in fluorescence intensity over time. For GFP-LC3B puncta that were present at the start of the time-lapse series, only those puncta that increased in fluorescence intensity and/or area with time were counted as AV biogenesis events. To quantify autophagosome transport, kymographs were generated in FIJI using the MultiKymograph plugin (line width 5) and analyzed in FIJI. Autophagic vesicles were classified as anterograde (net movement ≥ 10 μm toward the axon distal tip), retrograde (net movement ≥ 10 μm toward the cell body), or bidirectional/stationary (net movement < 10 μm). Time-lapse videos were referenced during kymograph analysis to ensure that extraneous neurites were not included in the data. Transport parameters were calculated in Microsoft Excel from the coordinates identified in FIJI.

To visualize lysosomes, LysoTracker (ThermoFisher) or Cresyl Violet was added to DIV2, untransfected DRGs. LysoTracker was added at 25 nM and incubated for 30 min at 37°C in a 5% CO_2_ incubator. Cell Mask Deep Red (ThermoFisher) was added for the last 5 min of the incubation. Cells were washed once with equilibrated complete F-12 prior to imaging. Cell Mask Deep Red and Cresyl Violet (1 μM) were added for 5 min and washed three times prior to imaging.

Z-stack micrographs were initially processed in FIJI into maximal projections. Maximal projection micrographs were segmented using Ilastik^70^. At least one image from each biological replicate was used to train Ilastik to identify lysosomes (LysoTracker or Cresyl Violet channels) and cell limits (Cell Mask channel). Training images were not used in subsequent data analysis. Images from all biological replicates for each channel were processed in batch mode by Ilastik to yield simple segmentation files. Segmented files were analyzed in FIJI using Analyze Particles.

### Immunoblotting

Brains of non-transgenic mice were dissected and subsequently homogenized and lysed. Brains were homogenized individually in RIPA buffer [1x PBS, 1% Triton X-100, 0.5% deoxycholate, 0.1% SDS, 1x complete protease inhibitor mixture (Roche), and 1x Halt™ protease and phosphatase inhibitor cocktail (ThermoFisher)]. Total protein in each lysate was determined by BCA assay (ThermoFisher Scientific) to ensure equal protein loading during western blot analysis.

All supernatants were analyzed by SDS-PAGE western blot, transferred onto FL PVDF membranes (MilliporeSigma), and visualized with fluorescent secondary antibodies (Li-Cor, ThermoFisher) using an Odyssey^®^ CLx imaging system (Li-Cor). See Key Resources Table for antibodies used. All western blots were analyzed with Image Studio (Li-Cor). Total protein was used as a loading control (REVERT^™^ Total Protein Stain, Li-Cor). The normalization factor is listed below each blot as a percent.

### Additional Methods

Figures were assembled in Adobe Illustrator. Prism 6, 8, and 9 (GraphPad) were used to plot graphs and perform statistical tests. Statistical tests are indicated in the text and figure legends.

## Key Resources Table

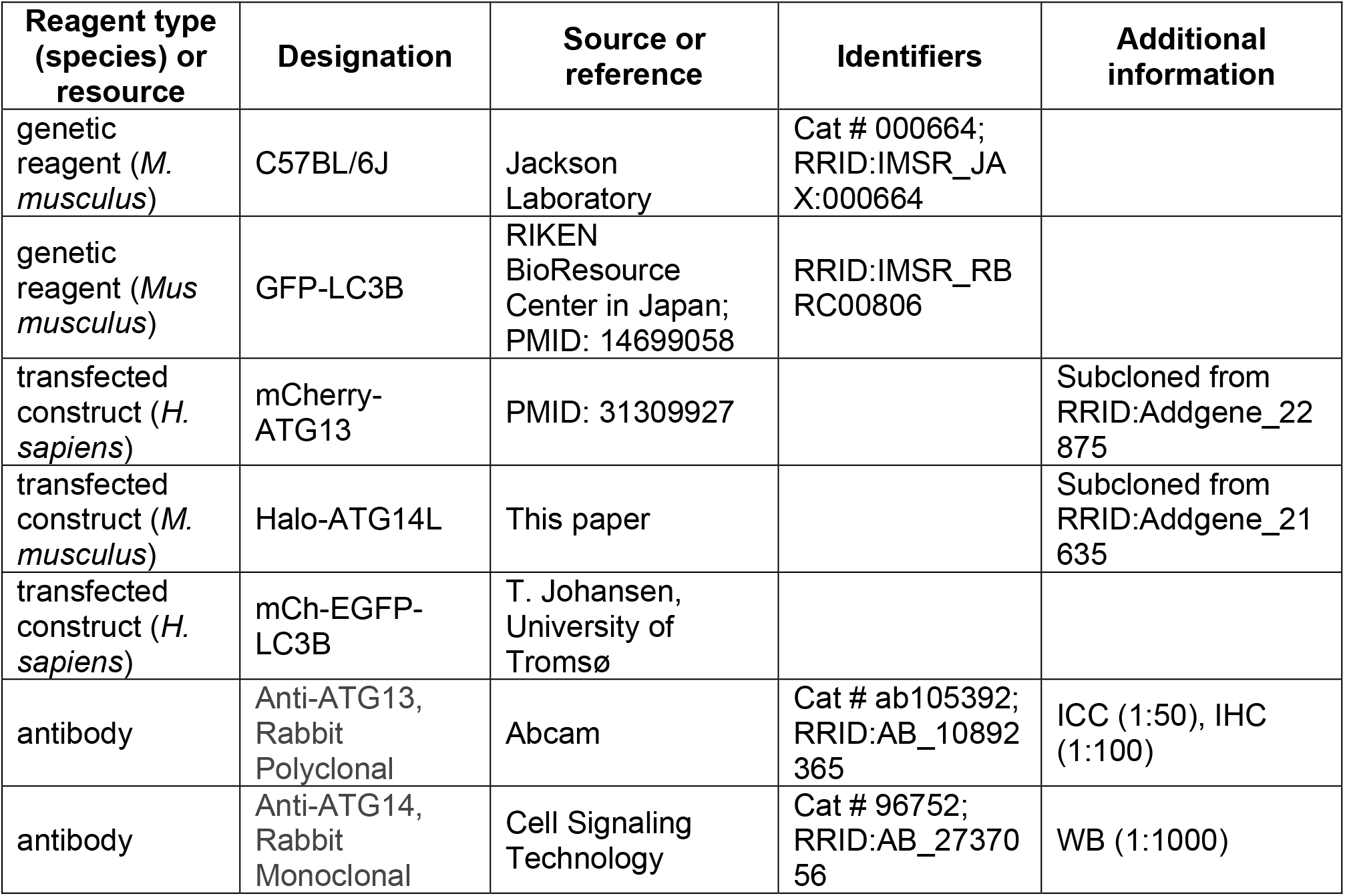
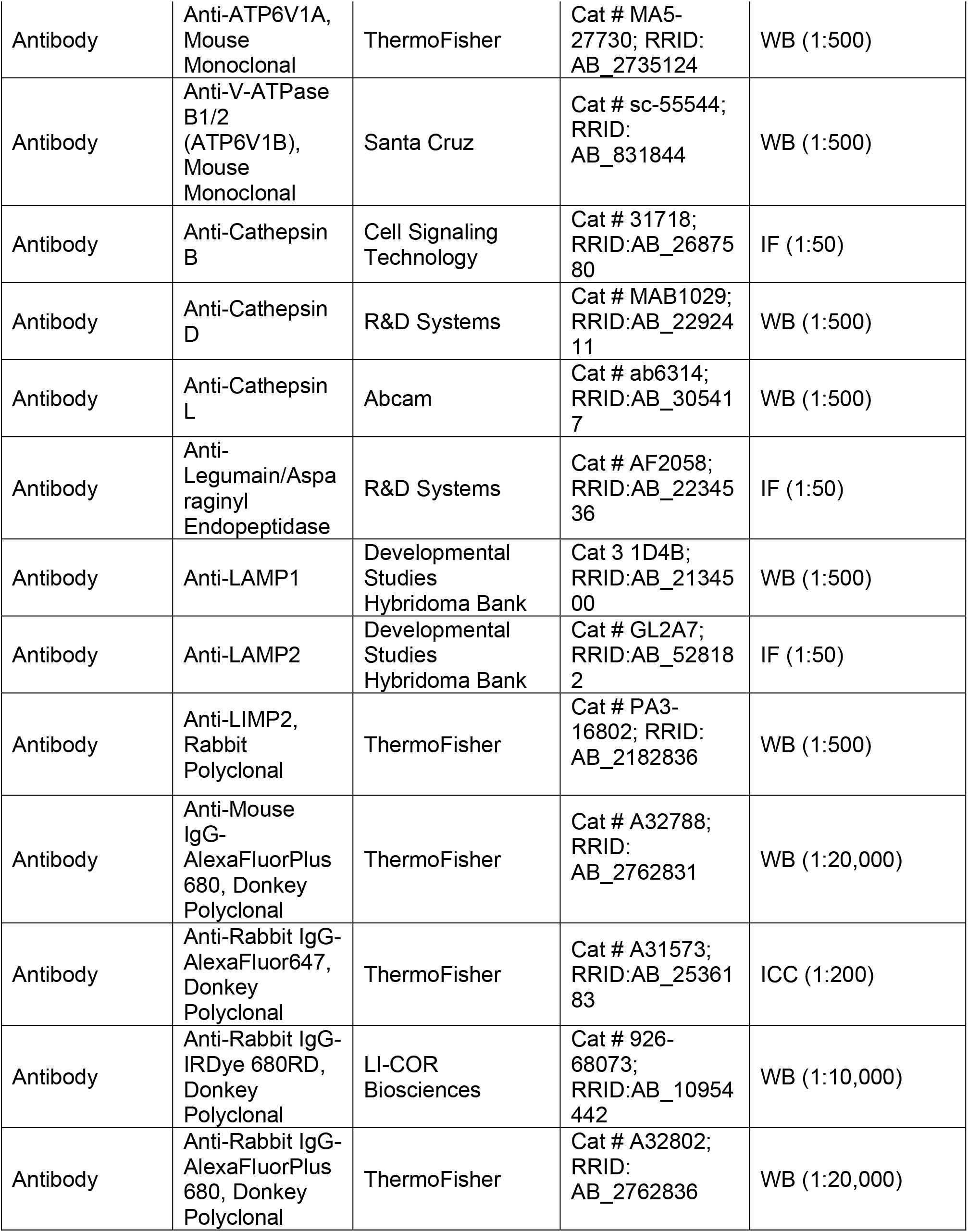
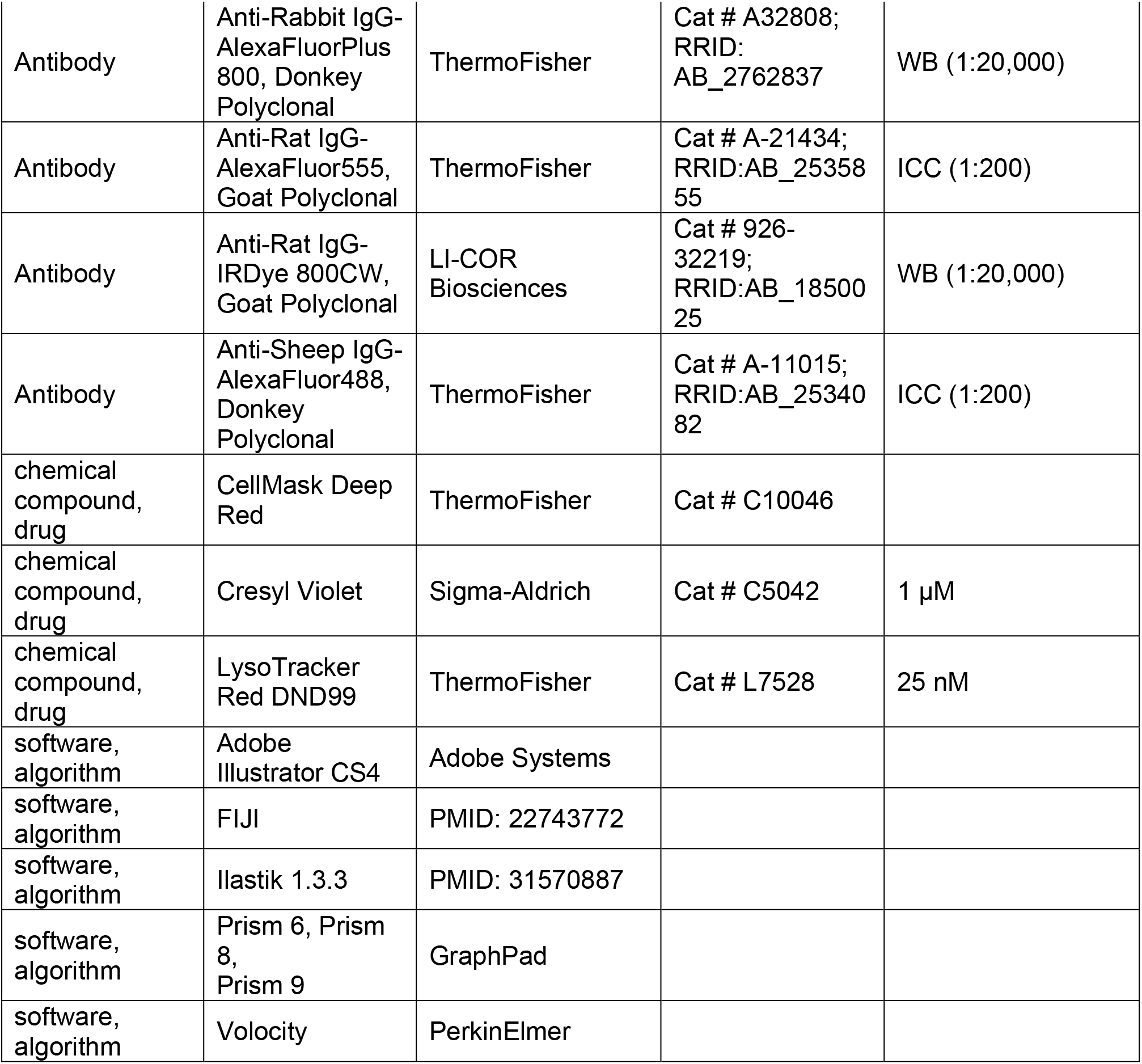

