## Supplemental Figures and Legends for "Aging Differentially Affects Axonal Autophagosome Formation and Maturation"

### SUPPLEMENTAL FIGURE LEGENDS

**Figure S1. Autophagy and lysosomal protein levels.**

(A) Immunoblots of brain lysates from young, young adult, and aged mice of both sexes (n=4 biological replicates shown for each age, blots were repeated on another set of 4 biological replicates for each age). Total protein was used as a loading control (normalization factor indicated below each blot as a percentage). 40 µg total protein was loaded in each lane. (B) Quantification of protein levels, normalized to total protein and represented as fold change relative to the mean for lysates from young (1 mo) mice. For individual data points, circles represent female, triangles represent male. *p < 0.05; **p < 0.01; ns, not significant by One-way ANOVA with Tukey's multiple comparisons test. (C) Immunoblots of brain lysates from young, young adult, and aged mice of both sexes (n=3 biological replicates shown for each age, blots were repeated on another set of 3 biological replicates for each age). Total protein was used as a loading control (normalization factor indicated below each blot as a percentage). 40 µg total protein was loaded in each lane. (C) Quantification of protein levels, normalized to total protein and represented as fold change relative to the mean for lysates from young (1 mo) mice. For individual data points, circles represent female, triangles represent male. *p < 0.05; ns, not significant by One-way ANOVA with Tukey's multiple comparisons test.

**Figure S2. Autophagic vesicle numbers in the distal and proximal axon.**

AV counts in the distal (top) and proximal (bottom) axon for immature AVs (yellow) and mature AVs (red). AVs are binned by net direction travelled and age.

**Figure S3. Average speed of immature and mature AVs does not considerably change with age in the distal neurite.**

(A) Cartoon of DRG neuron depicting time-lapse imaging of immature AVs in the distal neurite. (B-D) Quantification of the average speed of immature autophagic vesicles that moved in a net anterograde (B), net bidirectional or stationary (C) or net retrograde (D) direction in the distal neurite of DRG neurons from young (1 mo, light yellow), young adult (3 mo, yellow), and aged (16-17 mo, dark yellow) mice (mean ± SEM; n≥ 18 puncta from three biological replicates). ns, not significant by Kruskal-Wallis test with Dunn's multiple comparisons test. (E) Cartoon of DRG neuron depicting time-lapse imaging of mature AVs in the distal neurite. (F-H) Quantification of the average speed of mature autophagic vesicles that moved in a net anterograde (F), net bidirectional or stationary (G) or net retrograde (H) direction in the distal neurite of DRG neurons from young (light red), young adult (red), and aged (dark red) mice (mean ± SEM; n≥ 15 puncta from three biological replicates). *p < 0.05; ns, not significant by Kruskal-Wallis test with Dunn's multiple comparisons test.

**Figure S4. Pause time fraction of immature and mature AVs does not considerably change with age in the distal neurite.**

(A) Cartoon of DRG neuron depicting time-lapse imaging of immature AVs in the distal neurite. (B-D) Quantification of the pause time fraction of immature autophagic vesicles that moved in a net anterograde (B), net bidirectional or stationary (C) or net retrograde (D) direction in the distal neurite of DRG neurons from young (1 mo, light yellow), young adult (3 mo, yellow), and aged (16-17 mo, dark yellow) mice (mean ± SEM; n≥ 18 puncta from three biological replicates). ns, not significant by Kruskal-Wallis test with Dunn's multiple comparisons test. (E) Cartoon of DRG neuron depicting time-lapse imaging of mature AVs in the distal neurite. (F-H) Quantification of the pause time fraction of mature autophagic vesicles that moved in a net anterograde (F), net bidirectional or stationary (G) or net retrograde (H) direction in the distal neurite of DRG neurons from young (light red), young adult (red), and aged (dark red) mice (mean ± SEM; n≥ 15 puncta from three biological replicates). $^{**}p < 0.01$; ns, not significant by Kruskal-Wallis test with Dunn's multiple comparisons test.

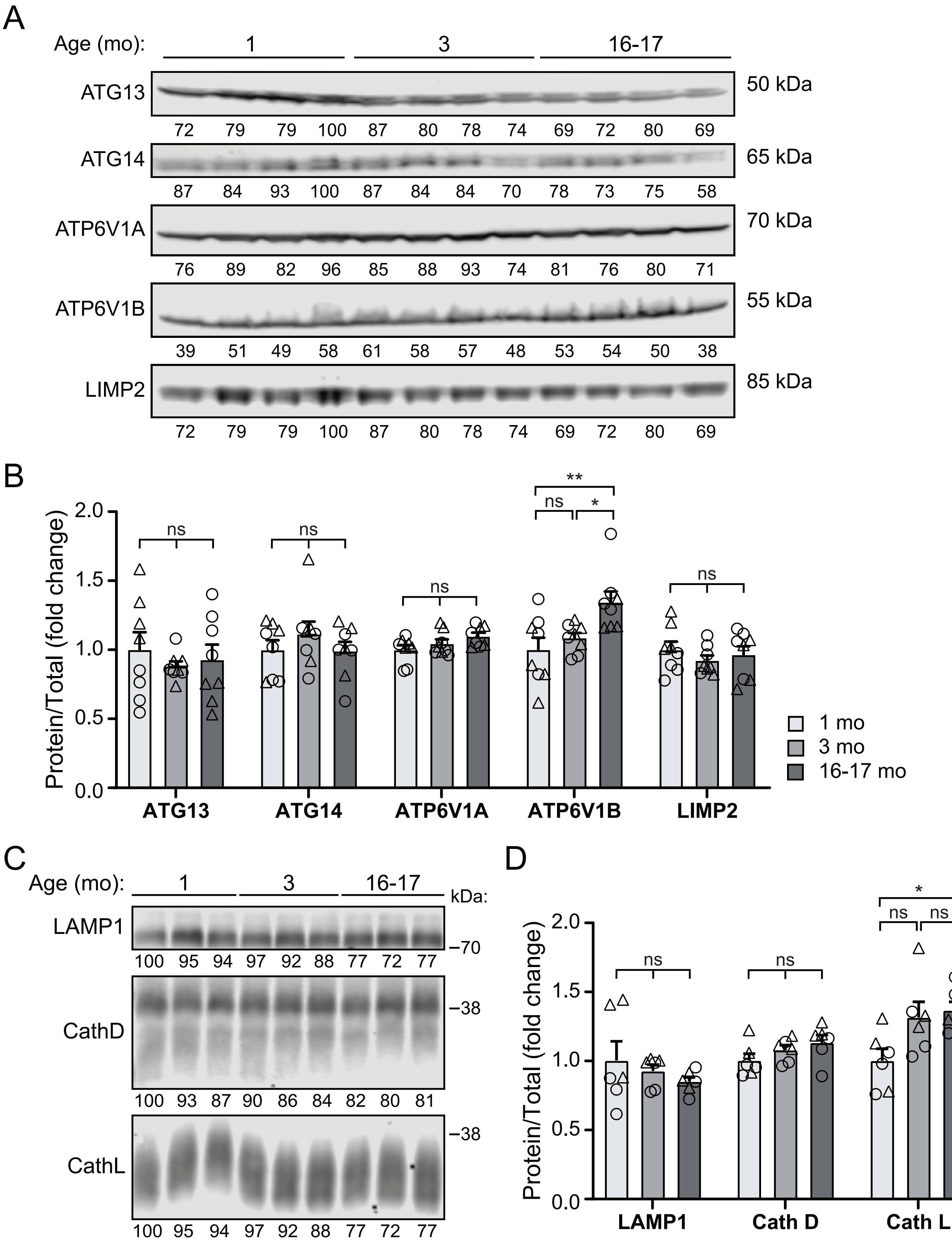
A
Age (mo):
1
3
16-17
ATG13
50 kDa
72 79 79 100 87 80 78 74 69 72 80 69
ATG14
65 kDa
87 84 93 100 87 84 84 70 78 73 75 58
ATP6V1A
70 kDa
76 89 82 96 85 88 93 74 81 76 80 71
ATP6V1B
55 kDa
39 51 49 58 61 58 57 48 53 54 50 38
LIMP2
85 kDa
72 79 79 100 87 80 78 74 69 72 80 69
B
Protein/Total (fold change)
2.0
1.5
1.0
0.5
0.0
ns
ns
ns
**
ns
*
ns
ATG13
ATG14
ATP6V1A
ATP6V1B
LIMP2
1 mo
3 mo
16-17 mo
C
Age (mo):
1
3
16-17
kDa:
LAMP1
–70
100 95 94 97 92 88 77 72 77
CathD
–38
100 93 87 90 86 84 82 80 81
CathL
–38
100 95 94 97 92 88 77 72 77
D
Protein/Total (fold change)
2.0
1.5
1.0
0.5
0.0
ns
ns
*
ns
ns
LAMP1
Cath D
Cath L

FIGURE S1

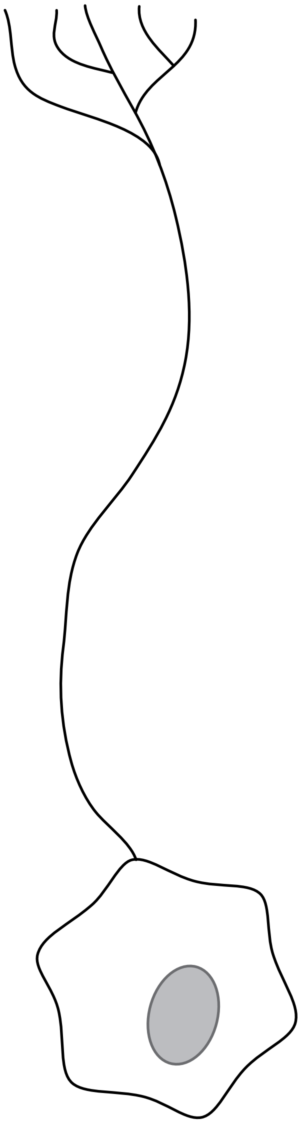

| Axon Location | AV Maturation | AV Net Direction | Age (mo) | # AVs |
|---|---|---|---|---|
| Distal | Immature | Anterograde | 1 | 18 |
| | | | 3 | 29 |
| | | | 16-17 | 20 |
| | | Bidirectional | 1 | 120 |
| | | | 3 | 73 |
| | | | 16-17 | 39 |
| | | Retrograde | 1 | 113 |
| | | | 3 | 63 |
| | | | 16-17 | 31 |
| | Mature | Anterograde | 1 | 36 |
| | | | 3 | 14 |
| | | | 16-17 | 15 |
| | | Bidirectional | 1 | 124 |
| | | | 3 | 62 |
| | | | 16-17 | 59 |
| | | Retrograde | 1 | 210 |
| | | | 3 | 176 |
| | | | 16-17 | 186 |

| Axon Location | AV Maturation | AV Net Direction | Age (mo) | # AVs |
|---|---|---|---|---|
| Proximal | Immature | Anterograde | 1 | 7 |
| | | | 3 | 4 |
| | | | 16-17 | 0 |
| | | Bidirectional | 1 | 38 |
| | | | 3 | 11 |
| | | | 16-17 | 14 |
| | | Retrograde | 1 | 92 |
| | | | 3 | 49 |
| | | | 16-17 | 21 |
| | Mature | Anterograde | 1 | 47 |
| | | | 3 | 15 |
| | | | 16-17 | 21 |
| | | Bidirectional | 1 | 130 |
| | | | 3 | 51 |
| | | | 16-17 | 54 |
| | | Retrograde | 1 | 298 |
| | | | 3 | 315 |
| | | | 16-17 | 242 |

FIGURE S2

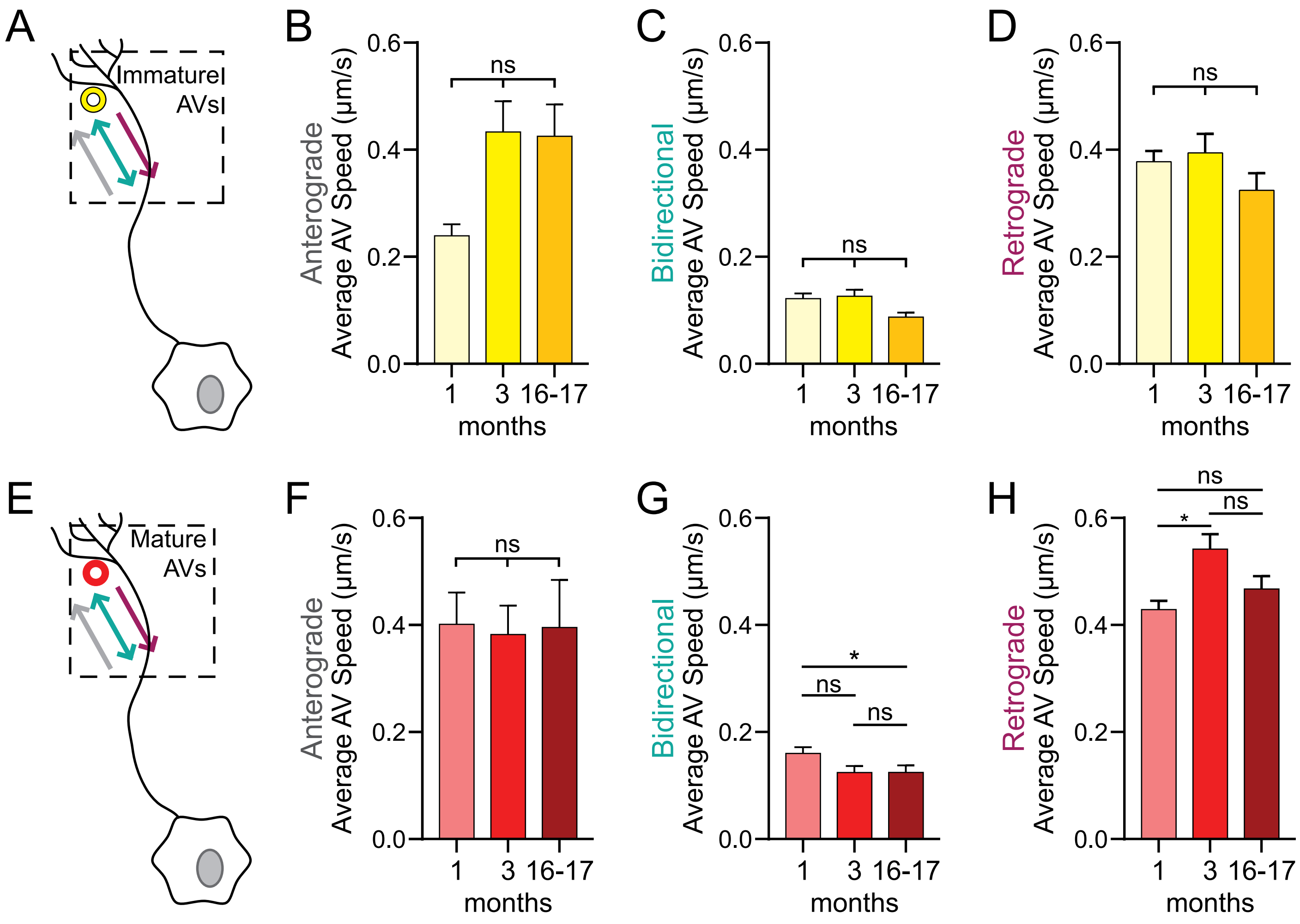
A
Immature
AVs
B
Anterograde
Average AV Speed (μm/s)
ns
0.0
0.2
0.4
0.6
1
3
16-17
months
C
Bidirectional
Average AV Speed (μm/s)
ns
D
Retrograde
Average AV Speed (μm/s)
ns
E
Mature
AVs
F
Anterograde
Average AV Speed (μm/s)
ns
G
Bidirectional
Average AV Speed (μm/s)
*
ns
ns
H
Retrograde
Average AV Speed (μm/s)
ns
*
ns

FIGURE S3

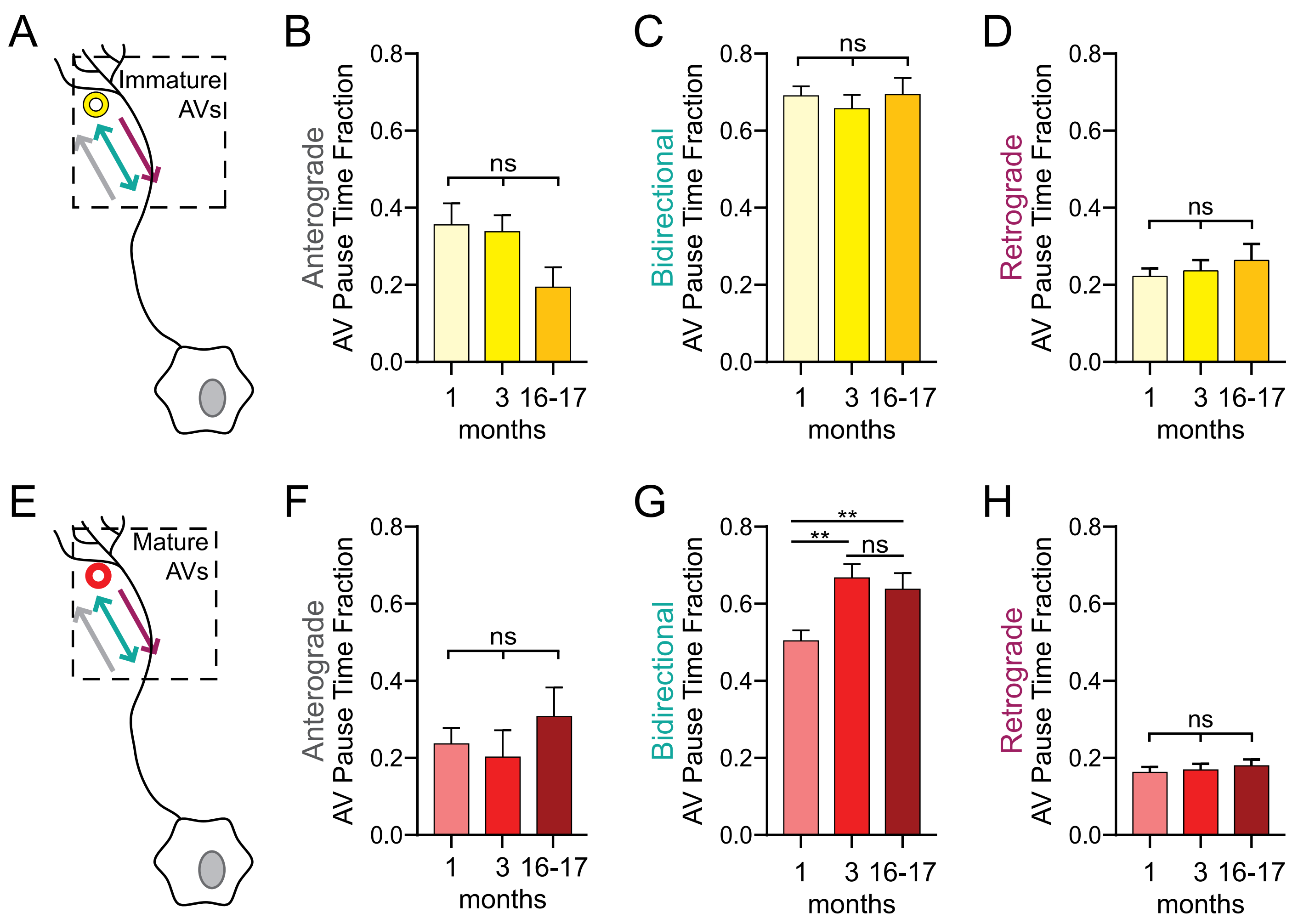

A
Immature
AVs
B
0.8
0.6
0.4
0.2
0.0
Anterograde
AV Pause Time Fraction
ns
1
3
16-17
months
C
Bidirectional
AV Pause Time Fraction
ns
D
Retrograde
AV Pause Time Fraction
ns
E
Mature
AVs
F
G
**
**
ns
H


FIGURE S4
